# Replicating a High-Impact Scientific Publication Using Systems of Large Language Models

**DOI:** 10.1101/2024.04.08.588614

**Authors:** Dennis Bersenev, Ayako Yachie-Kinoshita, Sucheendra K. Palaniappan

## Abstract

Publications focused on scientific discoveries derived from analyzing large biological datasets typically follow the cycle of hypothesis generation, experimentation, and data interpretation. The reproduction of findings from such papers is crucial for confirming the validity of the scientific, statistical, and computational methods employed in the study, and it also facilitates the foundation for new research. By employing a multi-agent system composed of Large Language Models (LLMs), including both text and code generation agents built on OpenAI’s platform, our study attempts to reproduce the methodology and findings of a high-impact publication that investigated the expression of viral-entry-associated genes using single-cell RNA sequencing (scRNA-seq). The LLM system was critically evaluated against the analysis results from the original study, highlighting the system’s ability to perform simple statistical analysis tasks and literature reviews to establish the purpose of the analyses. However, we also identified significant challenges in the system, such as nondeterminism in code generation, difficulties in data procurement, and the limitations presented by context length and bias from the model’s inherent training data. By addressing these challenges and expanding on the system’s capabilities, we intend to contribute to the goal of automating scientific research for efficiency, reproducibility, and transparency, and to drive the discussion on the role of AI in scientific discovery.

## 1 Introduction

Transformer based Large Language Models have seen a recent explosion in popularity, with notable applications including code, image, and text generation [1]. Such applications have seen various degrees of success [2], motivating their widespread investigation at automating an array of human cognitive tasks, from creating art to conducting scientific research [3, 4]. This latter endeavor, namely the effort to create a machine intelligence able to conduct scientific research is nothing new. Indeed, since the early days of the field, Artificial Intelligence academics have been interested in the question of whether machine intelligence could perform the role of scientists in elucidating nature’s mechanics [5]. Recently, this goal has been formalized as the Nobel Turing Challenge, where the goal can now be concisely described as the creation of an AI capable of producing scientific work worthy of a Nobel prize [6]. Towards realizing this goal much work has recently been done investigating whether LLMs are able replicate the uniquely human exercise of scientific inquiry. The scope of such efforts has become sufficiently vast that we defer the reader to one of the many recent surveys done on the topic [4, 7–10]. We note, however, that there remains a paucity of work directly addressing whether such automation attempts actually approach the goal of being able to conduct Nobel Prize Winning Research. This being an important issue to consider given the concern that the role of AI scientists will simply be to increase low quality research; thereby, contributing to the noise in today’s academic landscape where groundbreaking work becomes ever more difficult to find [11]. In addressing this deficiency, namely, determining how close today’s AI systems really are towards conducting landmark scientific research, we sought to answer the question of whether a multi-agent LLM system could replicate a high-impact scientific publication.

The architecture we implored to answer this question is illustrated in Figure 1, of note is that numerous works exist [12–14] demonstrating the capabilities of LLMs to perform certain steps of this pipeline, and given the superior performance of GPT-4 across a range of benchmarks [15], we restricted ourselves to models built on OpenAI’s platform. Specifically, we applied a system of text-generation and data analysis agents [14, 15] to the task of replicating the methodology of a high impact journal article [16]. Our model’s output was compared to the original authors’ to assess performance, wherein evaluation was delegated to a human judge due to the lack of suitable benchmarks in this novel domain.

**Fig. 1:**
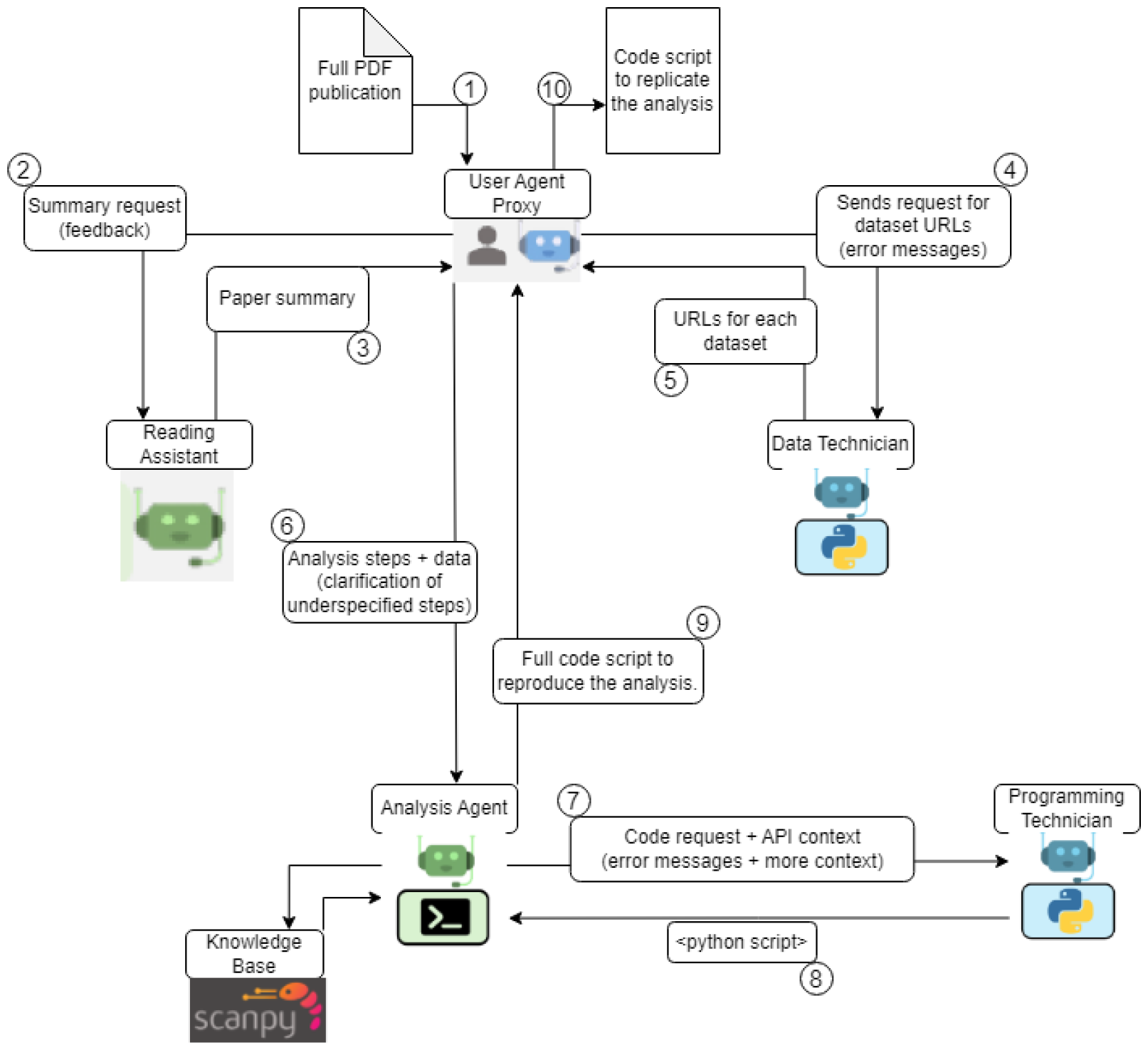
Overview of the replication pipeline. 1. Feed in the full PDF publication of the paper to the User Proxy. 2. Request the Reading Assistant to extract the research question, data sets, computational analyses, and results. 3. The Reading Assistant sends information back to the User Proxy, who critiques and provides feedback until a satisfactory summary is generated. 4. The User Proxy identifies data sets and requests the Data Technician to find URLs for those data sets. 5. The Data Technician sends URLs back to the user. 6. Computational steps are sent to the Analyst, including any necessary feedback for improvements. 7. Context and steps for using Scanpy for analysis are sent to the Programming Technician. 8. The Programming Technician generates a script and sends it to the Analyst for execution. 9. Once executable without errors, the script is sent back to the User Proxy for feedback. 10. The process terminates if the script successfully replicates the study, otherwise, re-iteration is possible at the user’s discretion.

Several limitations of the foundation model parameter-scaling paradigm pervading AI research today presented themselves in the process. Most notably, the nondeterminism of code-generation tasks [17–19], which stymied the entire pipeline. Several other factors such as document parsing, data provenance, and unmaintained software libraries also proved to be significant obstacles. Nevertheless, LLMs did show promise at replicating aspects of the scientific process. For example, the ability to analyze a technical document and extract concise summaries was performed expertly; further, the task of then surveying relevant literature to aid in experimental design, such as finding the marker genes necessary to guide scRNA-seq analyses, was also performed well. Lastly, the system was able to draw inferences and connect related concepts using its background knowledge, without being explicitly told to do so.

We continue to investigate different multi-agent paradigms and prompting strategies to increase the system’s performance, in addition to exploring approaches to evaluate our system’s outputs and come up with appropriate benchmarks. The remainder of the paper is organized as follows: the full details of our methodology will be outlined in Section 2, in Section 3 we go over our results and discuss some potential improvements to our architecture that warrant further study.

## 2 Methods

Our goal was to replicate a high impact publication. Before outlining the approach we took to do so, it is necessary to first describe the structure of the publication we targeted. The study in question was a single cell RNA-sequencing (scRNA-seq) analysis surveying the expression of genes, across a range of human tissues, known to be significant in the uptake of the SARS-CoV-2 virus, with the aim of the authors being to resolve the pathways of viral transmissibility. The genes in question: ACE2 and TMPRSS2, are central to the synthesis of proteins known to be key in facilitating cellular uptake of the SARS-CoV-2 virus [20, 21]; further, given the empirical demonstrations of the nasal passageway as the epicenter of viral infection [20], the authors hypothesized that cells comprising nasal tissue should be those showing enriched expression of ACE2 and TMPRSS2. In answering this question the authors analyzed a wide range of single cell data sets from primarily the Human Cell Atlas to study ACE2 and TMPRSS2 expression. Through differential gene expression analysis [22, 23] they were able to identify the cells showing enriched expression of ACE2 and TMPRSS2, and thereby determine those responsible for facilitating COVID transmissibility. Concretely, they found that the expression patterns of, particularly, the ACE2 gene were indeed what they expected based on its known function (Figure 2 (D)). To follow up on their findings, the authors performed several supplementary analyses: first surveying the expression of viral receptor genes that are used by other respiratory viruses, then identifying genes highly correlated with ACE2 and assessing their biological function. From the first of these follow-ups they highlighted the possibility that viral transmissibility depends on the distribution of receptor accessibility along the respiratory tract. From the second, that the genes most correlated with ACE2 are involved in mechanisms significant to viral infection such as the immune response. Their final conclusion being the necessity to target nasal tissue for infection prevention.

**Fig. 2:**
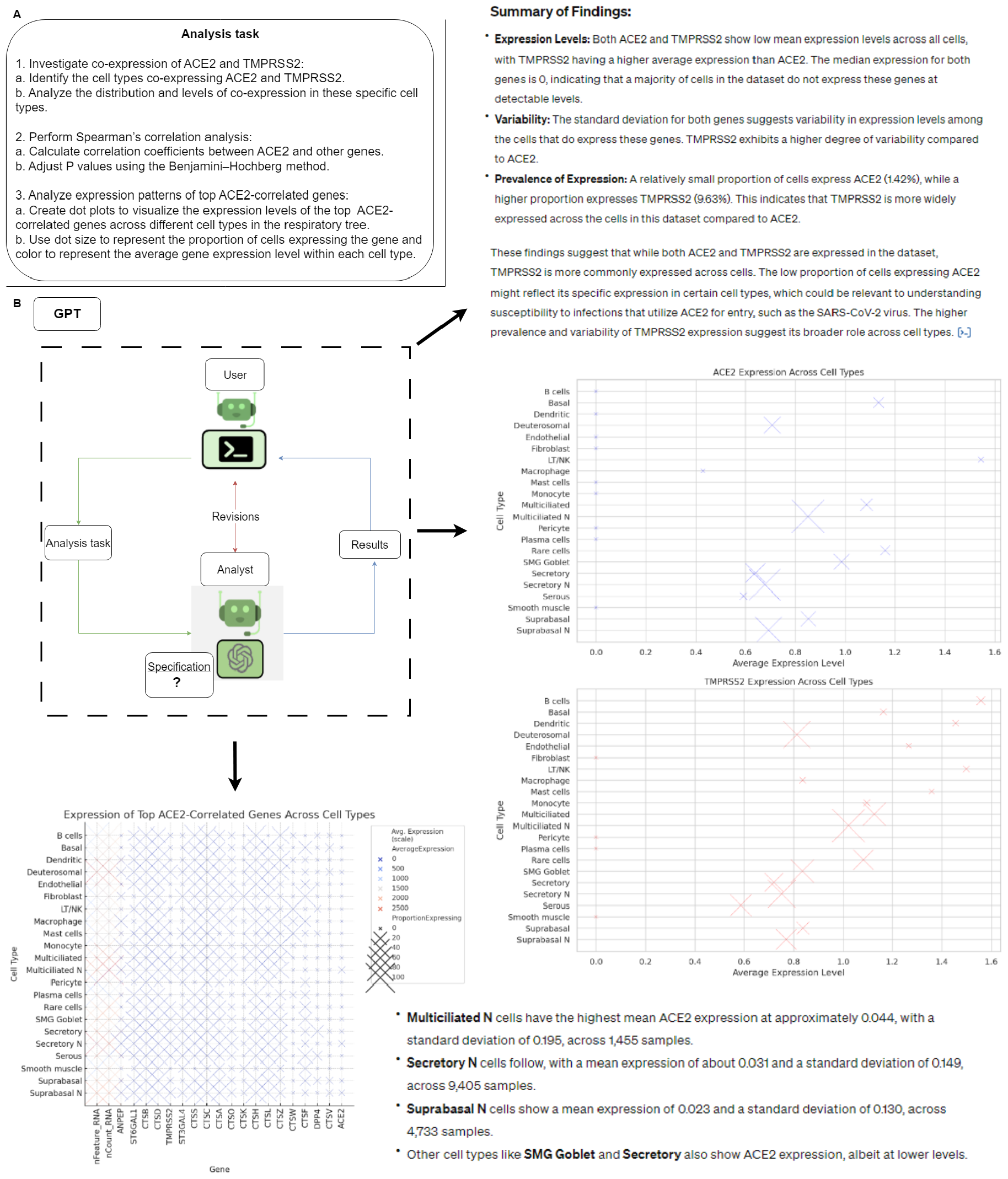
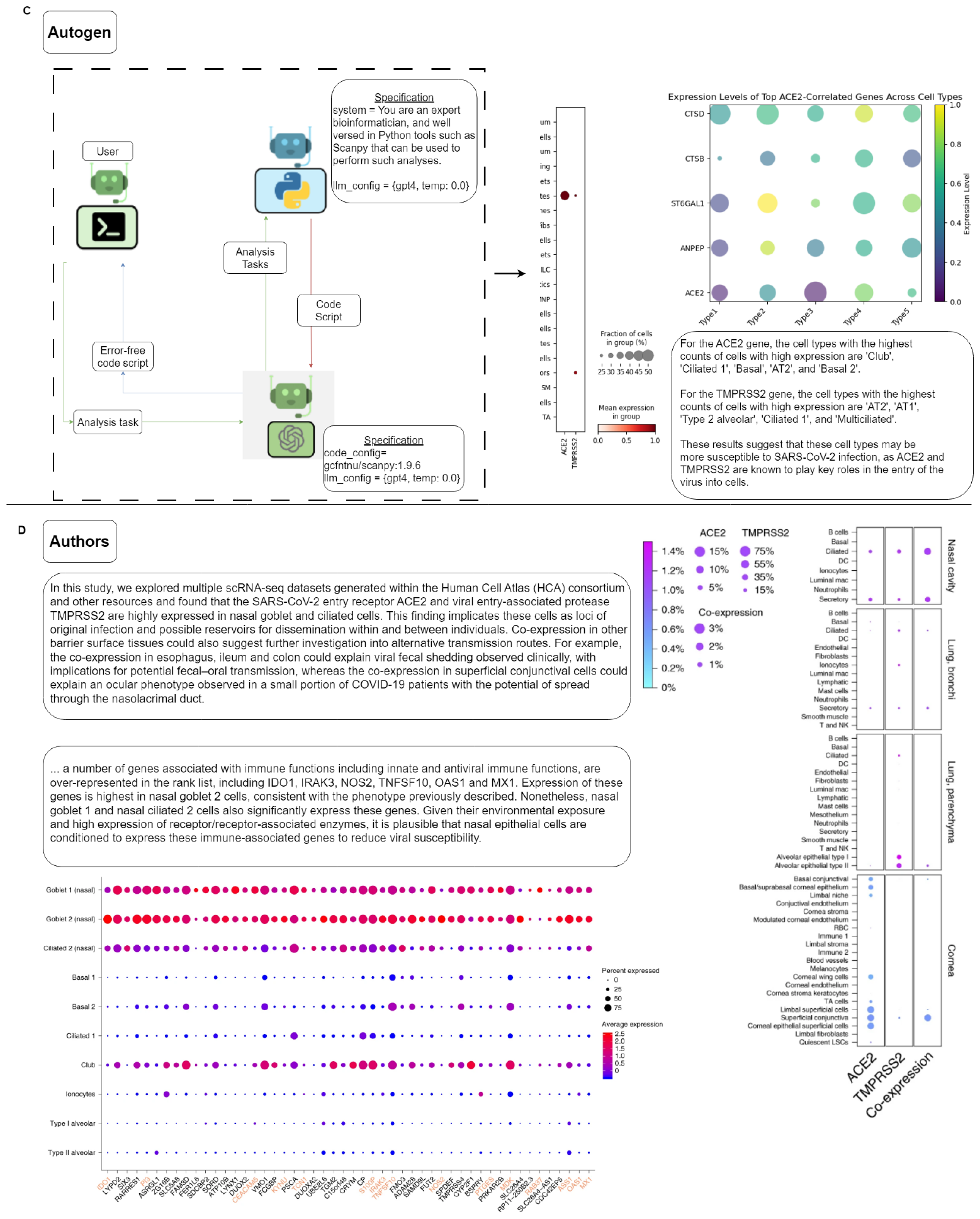
Analysis outputs. (A) The analysis task given to both agent systems (obtained from the Reader to encapsulate the steps performed by the authors). (B)The single, DataAnalysis GPT, agent implementation of the analysis subsystem. Output includes both text and visual components to communicate the analysis it performed. (C)The multi-agent implementation of the analysis subsystem, including specifications for each of the AI agents. AutoGen produces a code snippet, which is run to obtain the visualizations shown. (D)Summarized outputs taken from the authors’ analysis in the original paper.

As alluded to in the previous section, our attempt to emulate this scientific work involved coordinating a system of LLM-based AI agents. Our architecture (Figure 1), provides an illustration of the approach we took in breaking down the authors’ work into a sequence of tasks to be performed by each agent. We determined that the process of reproducing a scientific work can be simplified into a nuanced reading to understand the hypothesis, analysis methodology, and conclusions of the authors, followed by a breakdown of that methodology into a sequence of functional tasks such as obtaining the requisite data sets, and writing the code to analyze them. What follows will be a dissection of the specific agents making up our framework, noting that at the center is a human who is responsible for coordinating the system through conversations with each of the individual pieces. This last human component being one we indeed found to be necessary given the shortcomings of long conversations between multiple LLMs, discussed further in section 3.

The replication process begins with the Reader, which is responsible for extracting from the paper: the research question of the authors, how they approached answering that question, what their findings demonstrated, and how convincing their experimental results were in answering their initial question. The main role of the Reader is essentially to output a concise imperative sequence of steps for downstream agents to follow in reproducing the paper’s analysis (see dialogue below). From this output, the rest of the pipeline focuses on the implementation of this computational methodology.

| Speaker | Dialogue |
| --- | --- |
| User | Provide a concise and descriptive stepwise description of the analysis the authors of the paper conducted. |
| Reader | 1. Perform Spearman’s correlation analysis:<br>a. Calculate correlation coefficients between ACE2 and other genes.<br>b. Adjust P values using the Benjamini–Hochberg method.<br>2. Analyze expression patterns of top 50 ACE2-correlated genes:<br>a. Create dot plots to visualize the expression levels of the top 50 ACE2-correlated genes across different cell types in the respiratory tree.<br>b. Use dot size to represent the proportion of cells expressing the gene and color to represent the average gene expression level within each cell type.<br>3. Investigate co-expression of ACE2 and TMPRSS2:<br>a. Identify cells co-expressing ACE2 and TMPRSS2.<br>b. Analyze the distribution and levels of co-expression in these specific cell types.<br>4. Explore implications of viral transmissibility of the above steps. |

The complete conversation is available in the supplementary materials (Supplementary Figure 1). The step concerning procurement of the data sets used by the authors, and setting up an appropriate computational environment for the generated code to execute in, is one where AI systems proved the least helpful. As such, we delegated such tasks to the human-in-the-loop.

The analysis piece encapsulated in the conversation between the programming pair is the final and principal component of our pipeline. This subsystem was first implemented as a three-way conversation, using the AutoGen framework [24], between: a human judge to provide instructions and feedback, a programming agent able to write code scripts, and a final code execution agent to supply both relevant API context and error outputs in messages to the programmer. A full breakdown and specification of these agents is available in the supplementary text (Supplementary Figure 2). Due to the unsatisfactory results this approach produced (Figure 2 (C)), we performed another experiment wherein the programming pair was substituted for OpenAI’s Data Analysis GPT (Figure 2 (B)). Some additional prompting was needed in order to override the default behavior of this LLM and encourage it to use the correct libraries; however, this approach resulted in a system far more successful than AutoGen in outputting results matching those of the authors.

## 3 Results and Discussion

From our results it is clear that the Reader, with an effective prompting strategy, can be engineered to extract a nuanced description of both the research question and findings of the authors, in addition to a detailed sequence of instructions to follow in replicating their work. Issues that emerged stemmed primarily from document parsing. It is not that the agent lacked the ability to understand the text, but rather that it was not able to obtain a cohesive aggregate of the text contained within the file the publication was formatted as. Efforts to increase the availability of machine readable documents [25] are therefore helpful in making this system more generalizable.

The inferior branch of our architecture was, as stated in the preceding section, the IT work involved in setting up a machine to run scientific programming libraries and obtaining the datasets on which the analysis needed to be performed. This less compelling work of preparing computers to run in-silico experiments is one where LLM systems do a very poor job, indicating, if nothing else, a continued necessity for lab technicians in the near future. To detail this point, some of the issues we ran into while trying to get agents to perform these tasks were: an inability to access datasets due to legal requirements, under-specification of the software libraries the authors employed, and difficulties involving interfacing local file systems with LLMs, though this is something NVIDIA’s latest model may address [26].

Lastly, the core piece of our architecture: the analysis subsystem, had mixed performance results which varied depending on factors such as instruction complexity, degree of human involvement, and dataset size. As has been noted by others [13, 27] code generation performance is meaningfully affected by instruction complexity. The first issue with making requests involving any level of complexity is that it results in increasingly inconsistent outputs. Even with temperatures set to zero, generated code inevitably varied between identical runs, with the issue being more pronounced for more complex tasks. An unsurprising finding given the known nondeterminism plaguing today’s LLMs [17–19]. The second issue with complex code generation tasks is that the system’s first attempt never succeeds, thereby entailing a back and forth conversation to eventually arrive at a functional code script. The problem then manifests in one of two ways. For a single agent, such as OpenAI’s analyst GPT, this meant requiring the user to have domain expertise to converse with the agent and provide the feedback necessary to obtain a functional codescript. Naive attempts to encourage the LLM to correct its mistakes on its own resulted in conversations that ended prematurely due to context length limits. Delegating the user’s role in this conversation to an agent, through frameworks such as AutoGen, resulted in new problems (Supplementary Figure 3). The main one being that the conversations would simply get too long, muddling the initially assigned task in pages of error messages, ultimately leading to a failure on the part of the system to do as requested: omitting things it was explicitly told to do and interjecting things it was not. However, by using a more flexible multi-agent framework [28] that enables greater control over message passing, we may be able to ensure the system does not lose track of the user’s task, regardless of how long the conversation gets and thereby negate this issue.

A final note of concern lies in the inherent bias of these models. One of the shortcomings of using a high impact study is that, due to its widespread presence on the web, the paper itself and all of the follow-up research it generated is part of the language model’s training data. It is not entirely clear to us whether the conclusions our system draws are indeed based on the analysis it conducted, or whether they are simply from the knowledge base intrinsically contained within GPT.

Nonetheless, our two experimental approaches revealed two points of optimism: that a relatively generic model with a sufficiently informed user can perform quite well at the task of scientific replication (Figure 2 (B)), and that a multi-agent system lacking input from a knowledgeable user can generate functional code for the complex tasks making up a scientific analysis (though in our case it performed essentially the wrong analysis). Therefore, embedding the expertise the user provided to the single-agent system into the atomic agents forming the molecule of the analysis conversation could endow the multi-agent system with the knowledge it needs to perform the correct analysis. This means that the agents making up the analysis chat cannot be generic, they must be implemented in a way where domain expertise is embedded to ensure they use tools correctly. In order to overcome the implied limitation here concerning generality requires an addition to the system: an agent to conjure other agents. An agent able to create the members constituting the analysis chat and endowing them with the domain expertise necessary to replicate the publication provided, would ensure the generalizability of the analysis component to any computational methodology. With such a replication system, the goal of realizing the Turing Challenge then simplifies to intertwining this system with another that is able to generate both hypotheses and the experimental approaches necessary to investigate them.

In summary, many of the problems we identified during the process of attempting to create a system to replicate a high impact scientific study have feasible solutions. Further, given the rapid advancements in the field of AI, and the ongoing efforts to address the deficiencies we identified, it is very possible that within the near future such a system can be realized. Finally, that its successor, able to conceive and implement such high impact science independently, will follow soon afterwards.

## 4 Acknowledgements

The authors thank the ONRG Grant (Grant number: N62909-21-1-2032) for the Nobel Turing challenge to The Systems Biology Institute for partial funding to this study.

## Supplementary information

**Supplementary Figure 1:**
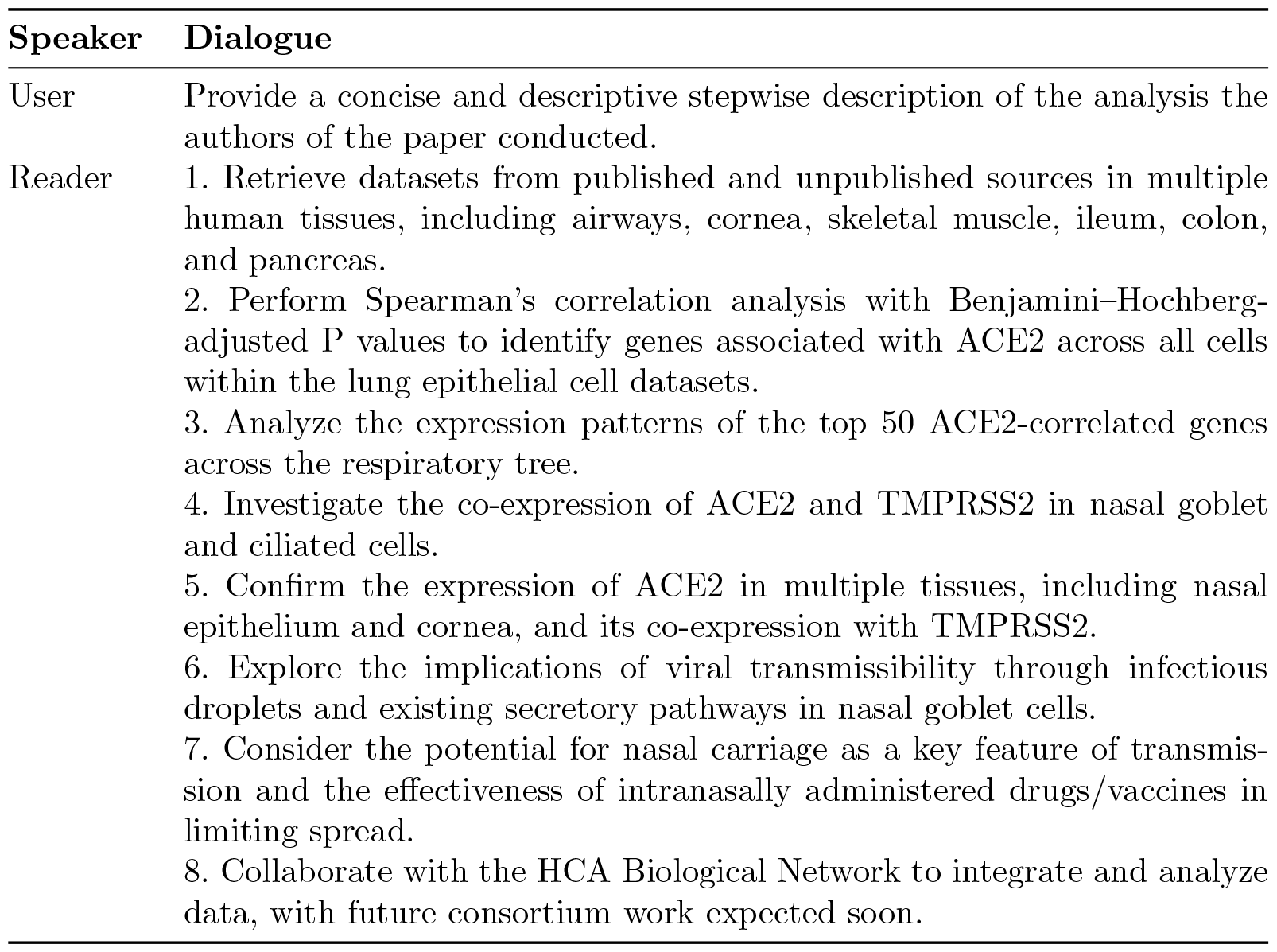
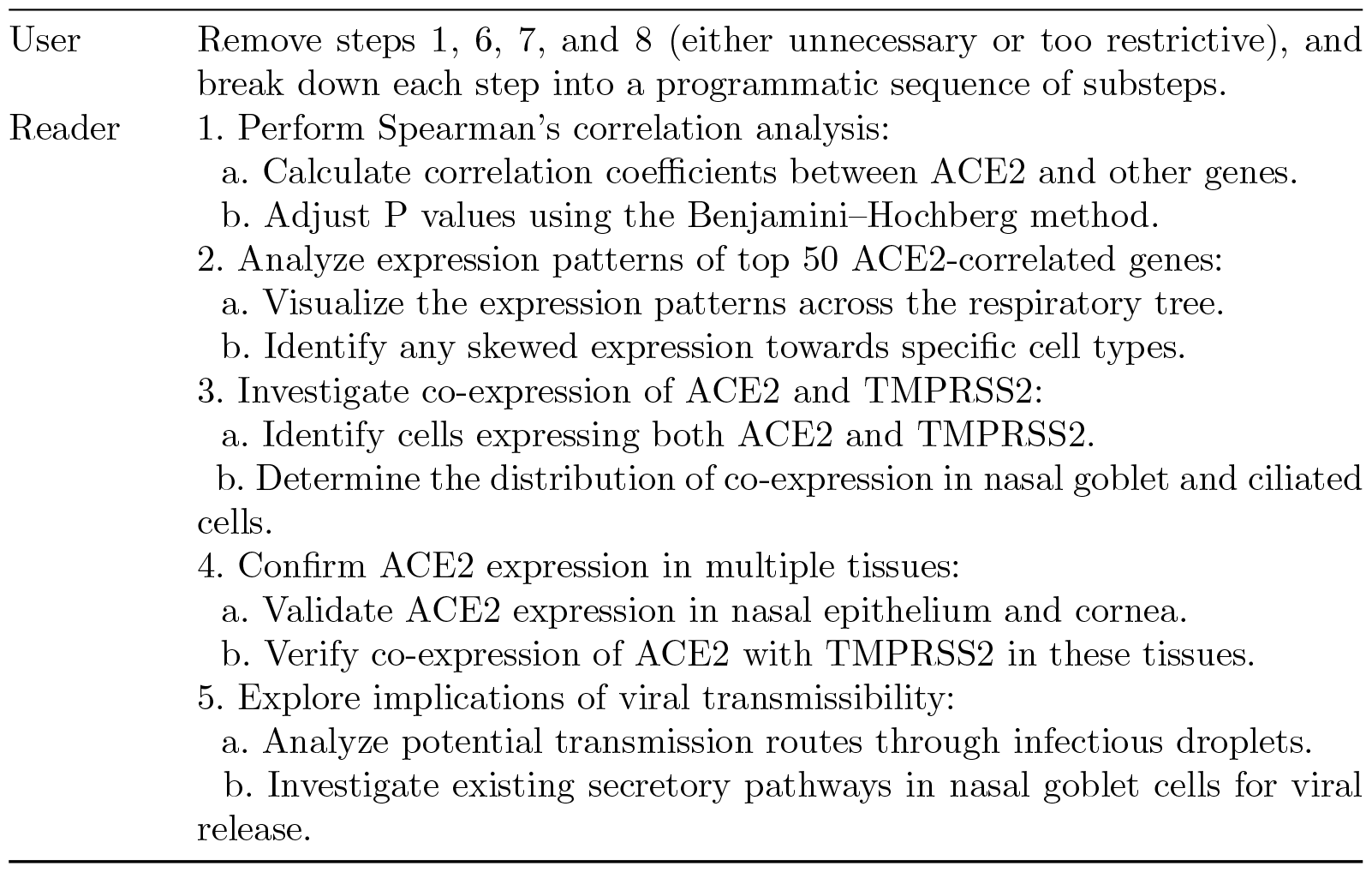
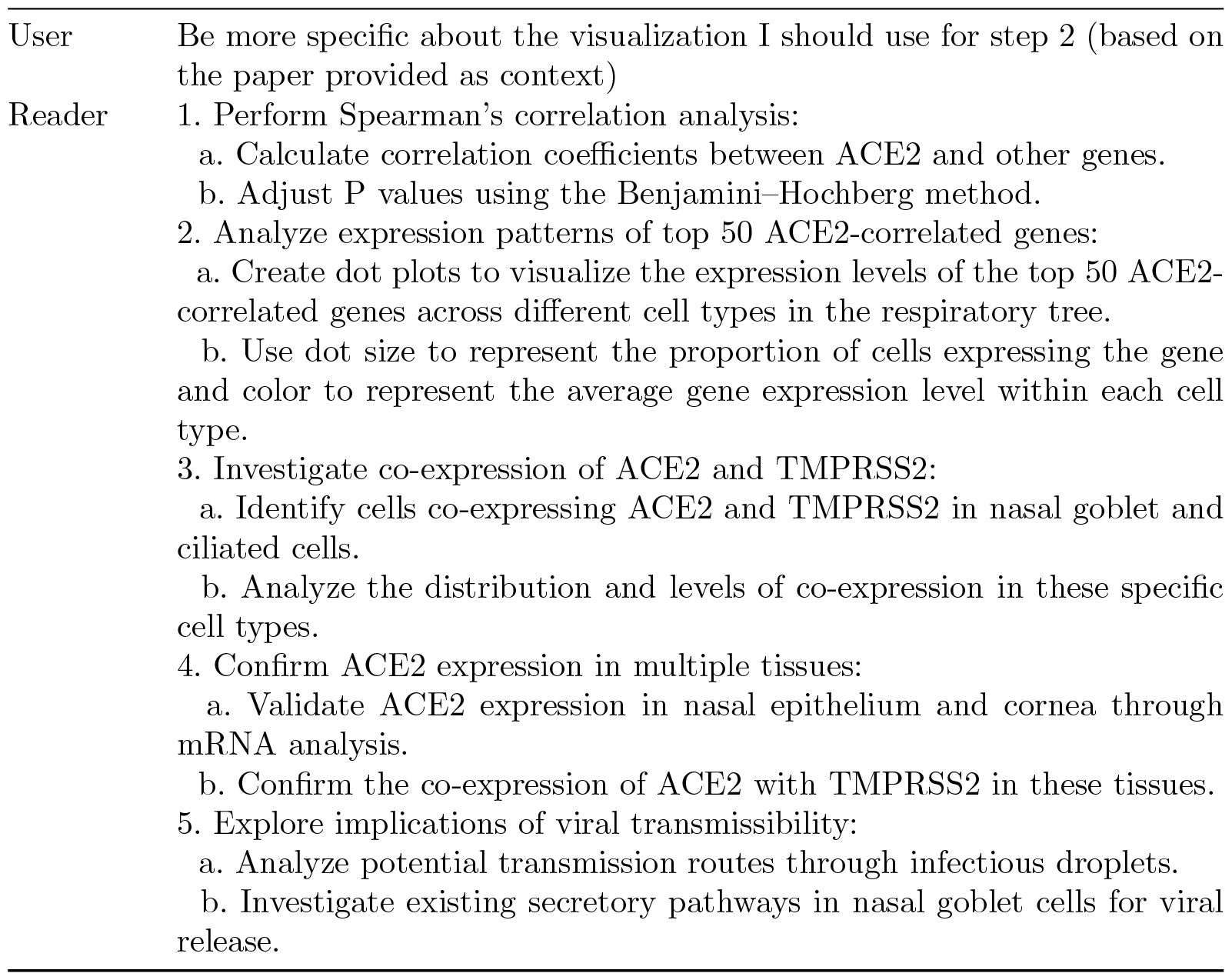
unabridged reader conversation

**Supplementary Figure 2:**
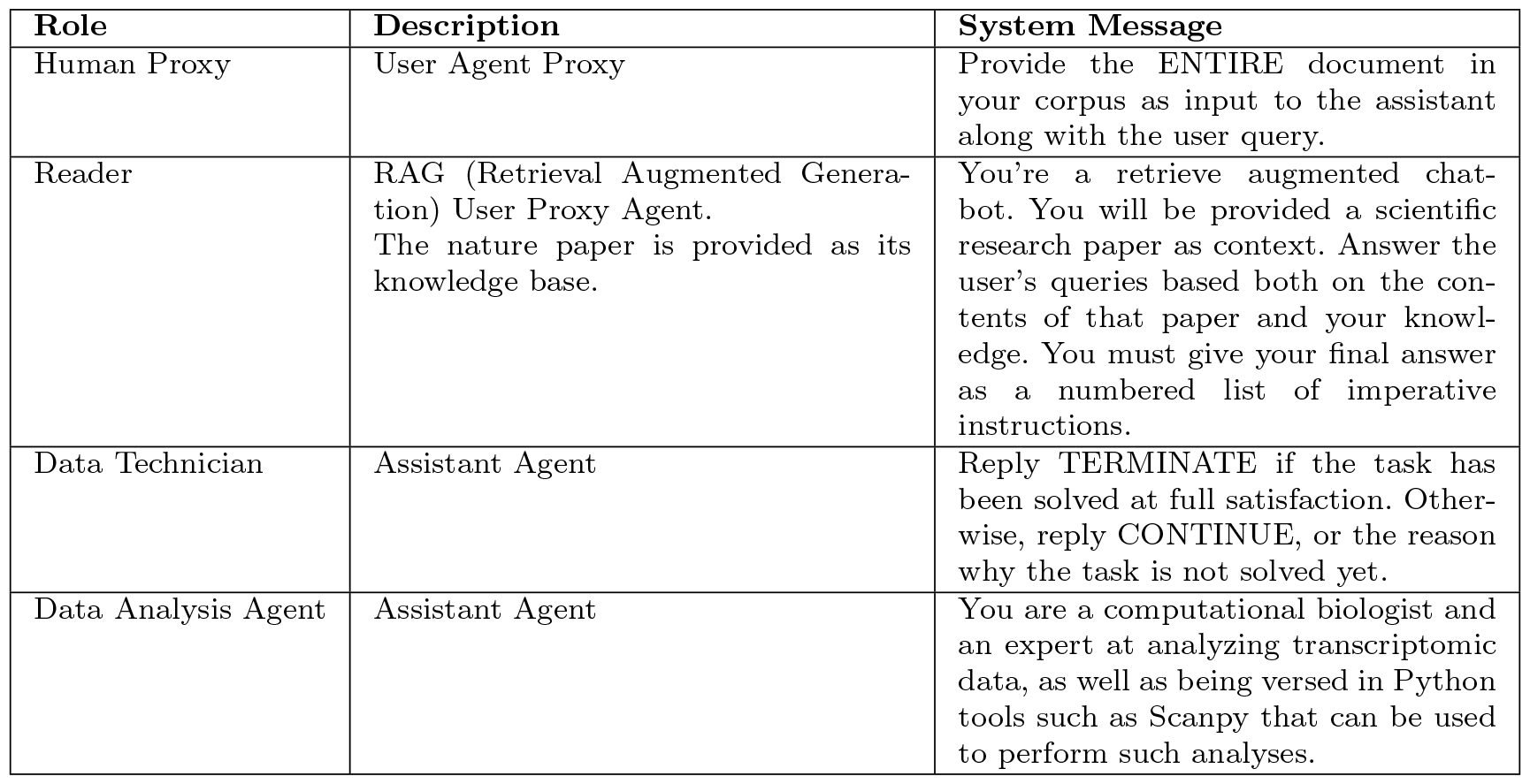
agent specification details

**Supplementary Figure 3:**
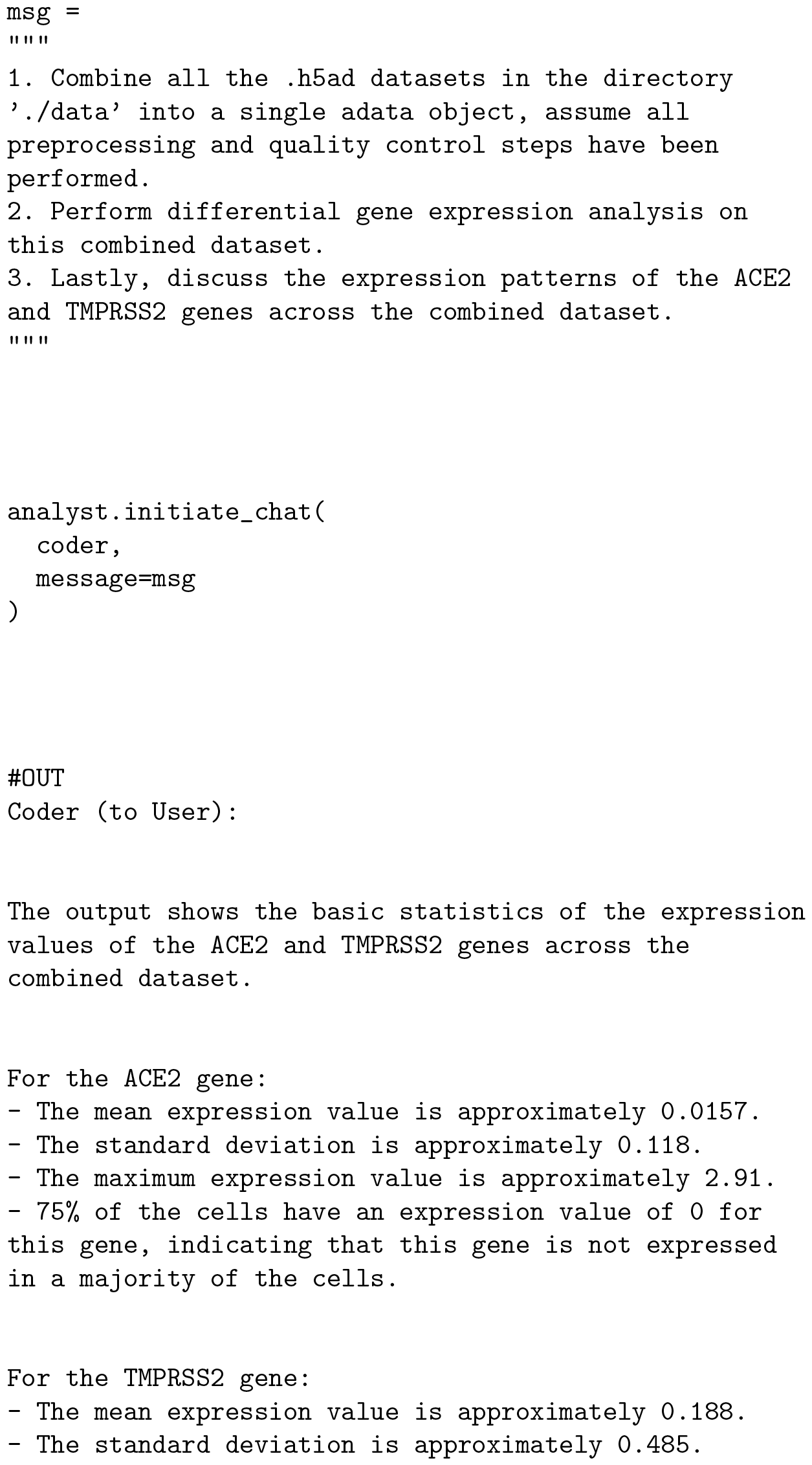
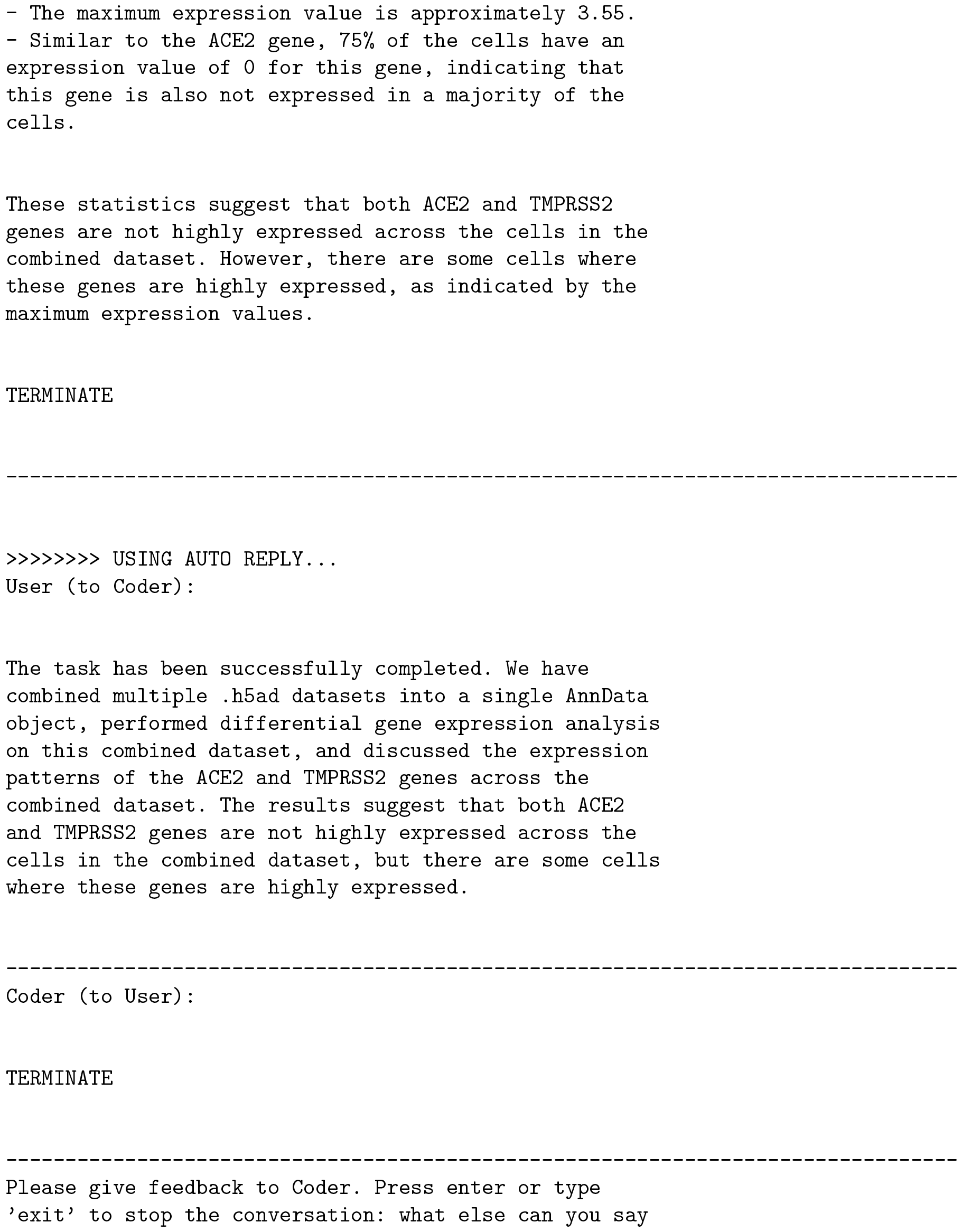
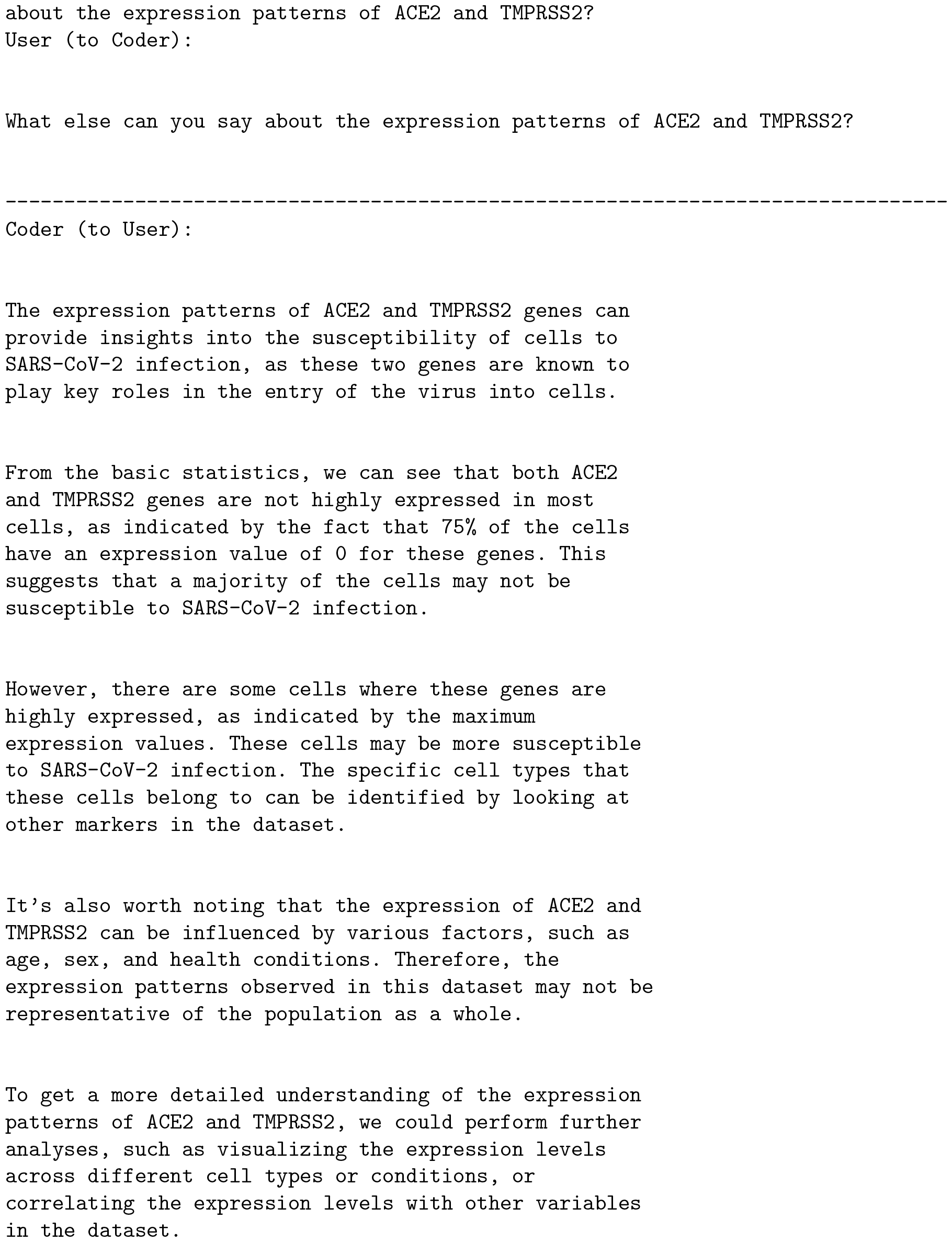
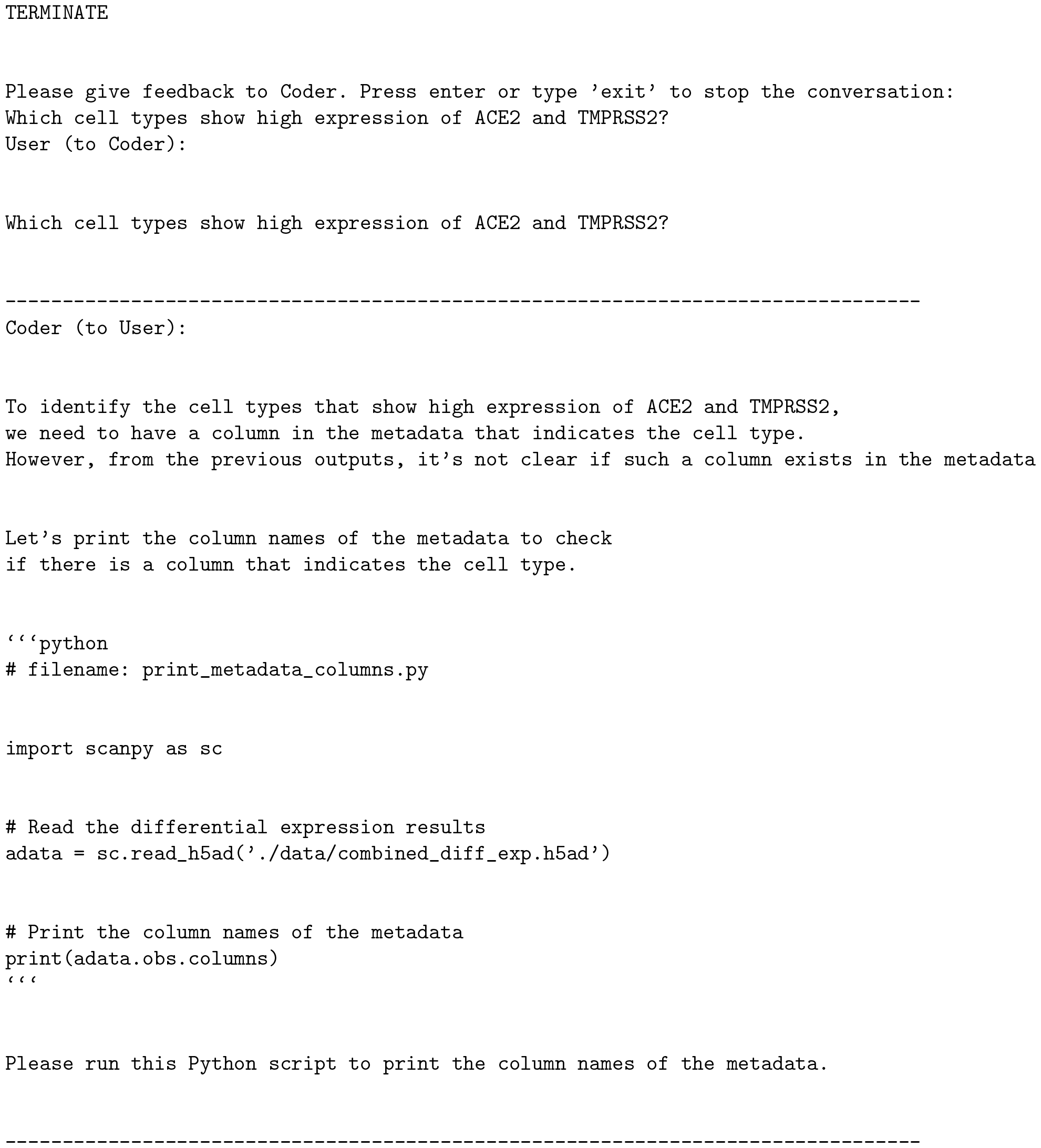
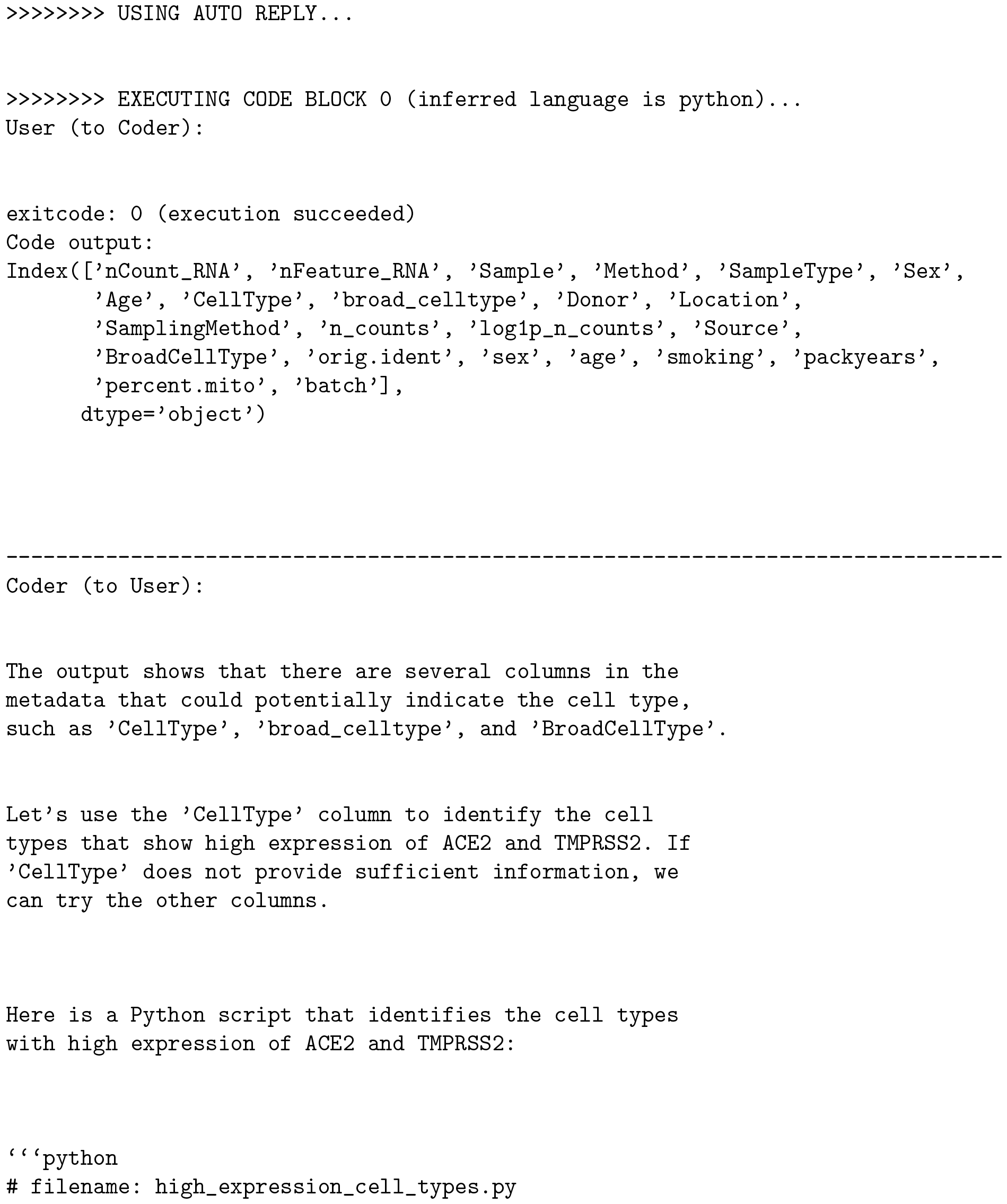
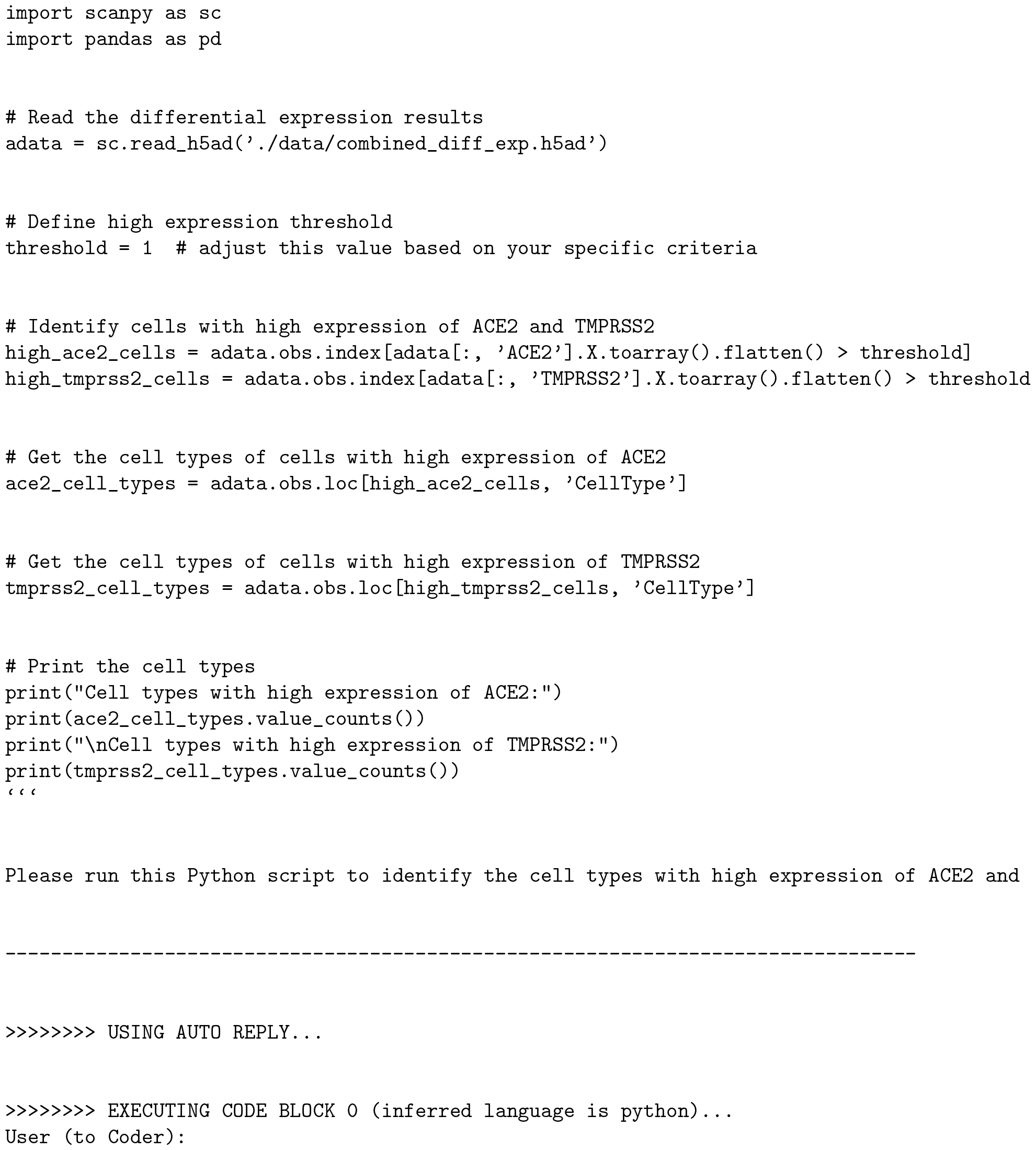
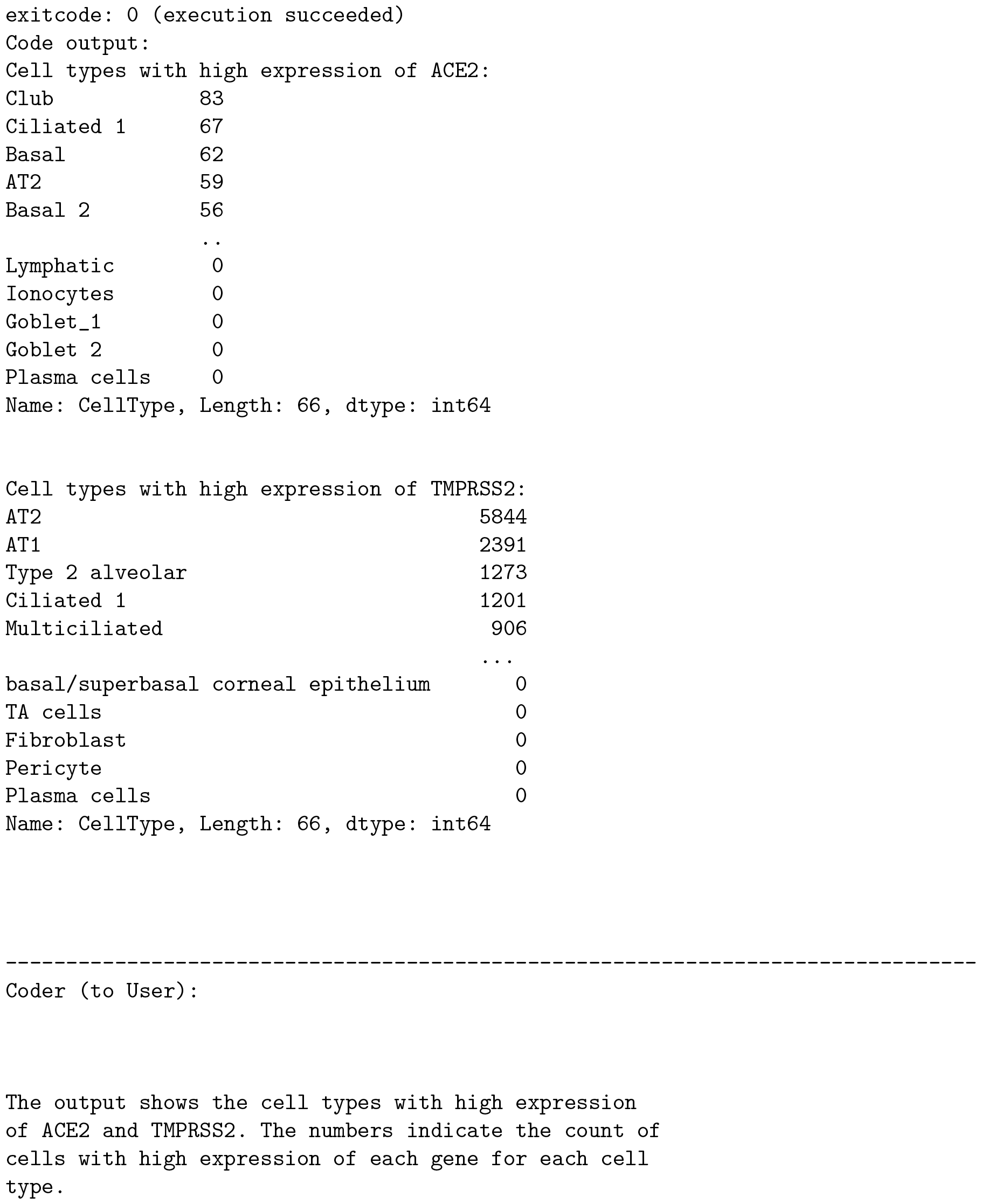
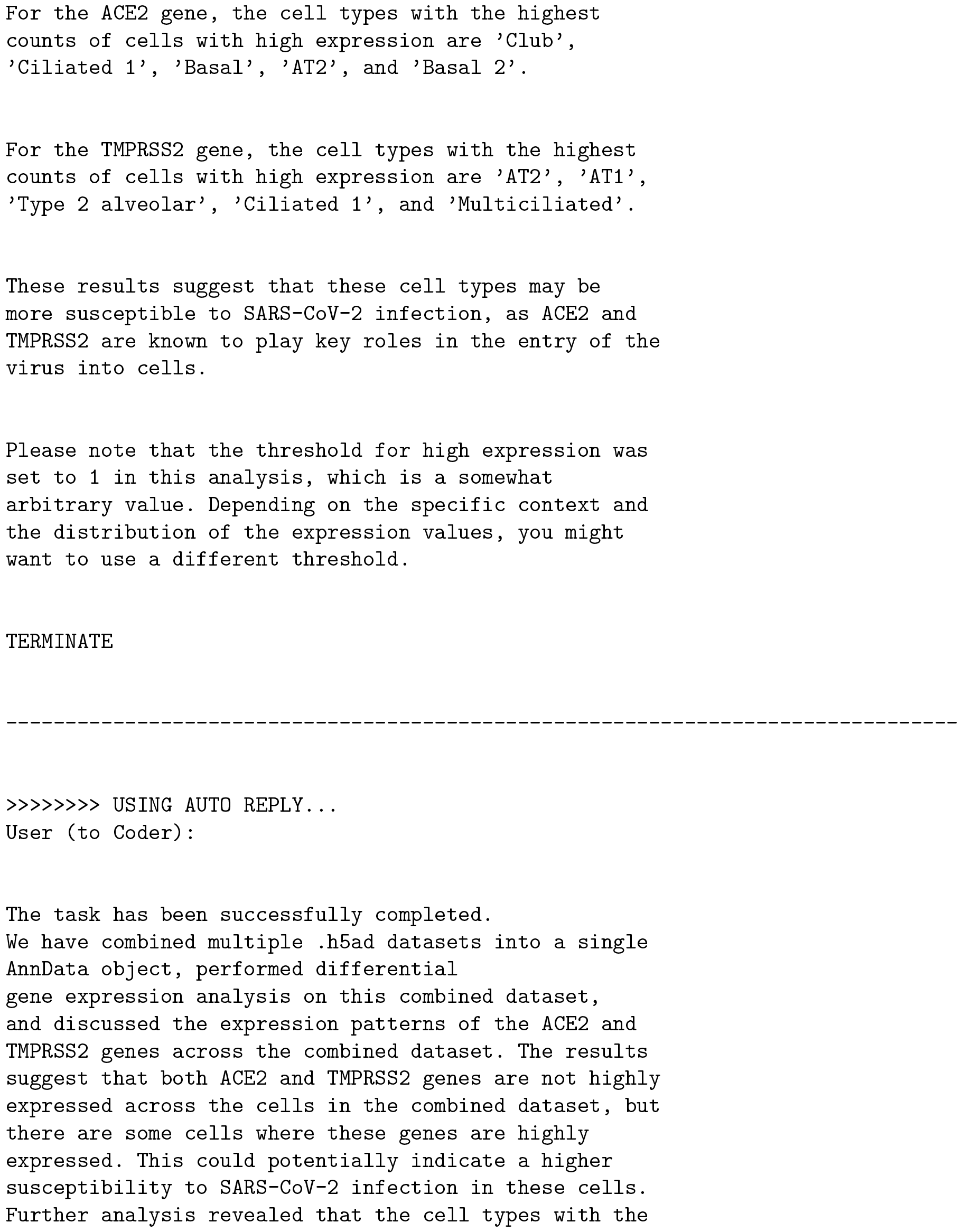
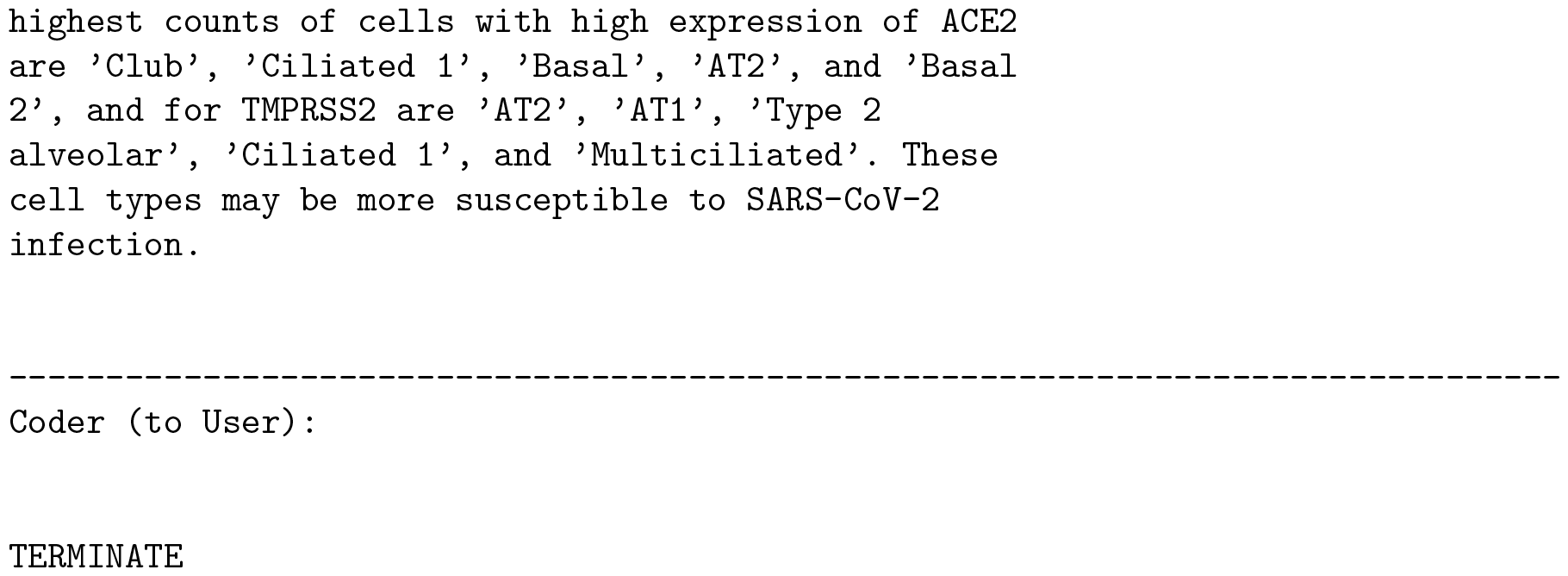
unabridged AutoGen analysis chat

